# Sleep-drive reprograms clock neuronal identity through CREB-binding protein induced PDFR expression

**DOI:** 10.1101/2019.12.12.874628

**Authors:** MK Klose, PJ Shaw

## Abstract

Neuronal circuits can be re-modeled by Hebbian plasticity, synaptic scaling and, under some circumstances, activity-dependent respecification of cell-surface receptors. Although the impact of sleep on Hebbian plasticity and synaptic scaling are well studied, sleep’s role in receptor respecification remains unclear. We demonstrate that high sleep-pressure quickly reprograms the *Drosophila* wake-promoting large-ventrolateral clock-neurons to express the Pigment Dispersing Factor receptor. The addition of this signaling input into the circuit is associated with increased waking and early mating success. The respecification of Pigment Dispersing Factor receptor in both young and adult large ventrolateral neurons requires two dopamine receptors and activation of the transcriptional regulator *nejire* (CREB-binding protein). These data identify receptor-respecification as an important mechanism to sculpt circuit function to match sleep levels with demand.

## Introduction

The expression of <u>specified</u> transmitters and receptors in maturing neurons can change prior to neuronal fate determination. As receptor expression profiles change, input sensitivities are altered and circuits are functionally remodeled^1^. Activity-dependent respecification of receptors can also occur in adult neurons in response to sustained increases in sensory and motor activity^2,3^ Thus, receptor respecification is a form of plasticity that, like Hebbian and homeostatic plasticity, may be employed to alter circuit function in response to changing environmental demands^3^. In mammals, birds, flies, fish and worms sleep circuitry is plastic and can change through developmental and in response to environmental factors (e.g. starvation, predation risk, mating status)^4–12^. Surprisingly, it remains unknown whether receptor respecification plays a role in modulating sleep-plasticity.

In *Drosophila*, the ventrolateral clock neurons (LNvs) regulate sleep, sleep-plasticity, light-arousal and many other clock influenced behaviors^13–17^. Much is known about the plasticity of neuropeptide expression levels in regulating circuit function in these neurons^18^, while much less is known about how receptor plasticity may contribute. In adult flies, the large LNvs (lLNvs) release the wake promoting peptide, *pigment dispersing factor (Pdf)*^19^ but do not endogenously express functional *Pdf receptor (Pdfr)* themselves^20^. However, we report here that functional PDFR is present in the lLNvs for the first ~48 h after eclosion, when sleep-drive is highest. Gain and loss of function experiments reveal that in young flies, PDFR expression is associated with increased waking and early mating success. Importantly, the PDFR can be re-established in adult lLNvs through prolonged sleep disruption. This respecification of PDFR in both young and adult lLNvs requires two G-protein coupled receptors sensitive to dopamine as well as activation of *nejire* (CREB-binding protein). These data identify receptor-respecification as an important mechanism to sculpt circuit function to match sleep need with environmental demands.

### PDFR is expressed in ILNvs in young flies

Sleep is highest in young animals during a critical period of brain development when neuronal plasticity is high^11,21^. As previously described in flies, sleep is highest during the first 48 h after eclosion (Day 0, Day1) and then reaches stable mature adult levels by day ~3 (Fig. 1a,b). The increased sleep observed during these ~48 h is important for the development of circuits that maintain adaptive behavior into adulthood^22,23^. How neurons in sleep circuitry change during this period has not been explored. The lLNvs promote waking behavior through both dopamine (DA) and octopamine (Oa) signaling (19, 22-24), thus we hypothesized one or both of these pathways might be downregulated during this early developmental period of high sleep. To test this hypothesis we used live-brain imaging in lLNv neurons expressing the reporter Epac1-camps to define cAMP response properties^16,20,24,25^. Contrary to our hypothesis, neither DA- or Oa-induced cAMP responses changed as the flies matured (Fig. 1c,d and Supplementary Fig. 1a,b). Interestingly, we did observe PDF-induced cAMP responses in lLNv neurons in the first 48 hours of adulthood (Figure 1 e,f), while they were predominantly absent in mature adult lLNvs, consistent with previous reports^20,24^. To determine if this transient PDF sensitivity is regulated at the receptor level, expression of the PDFR was examined directly using *Pdfr-myc*, a tagged receptor genetic construct under the natural PDF promoter^26^. As anticipated, detection of MYC antibody staining is high on day 0 and not detectable on day 5 of adulthood (Fig. 1 g) revealing transient expression of the receptor. Finally, we examined an adjacent group of clock neurons, the small ventrolateral neurons (sLNv)^15^. Responses to PDF in sLNvs are present at the beginning of adulthood, and then decrease in amplitude over the first ~48 hrs of adulthood. In contrast to the lLNvs, sensitivity to PDF in the sLNvs persists into mature adulthood (Supplementary Fig. 1c,d). Together these data indicate that the PDFR is transiently expressed in wake-promoting lLNvs in young flies when sleep-drive is high and is reduced or absent in mature adults.

**Fig. 1.**
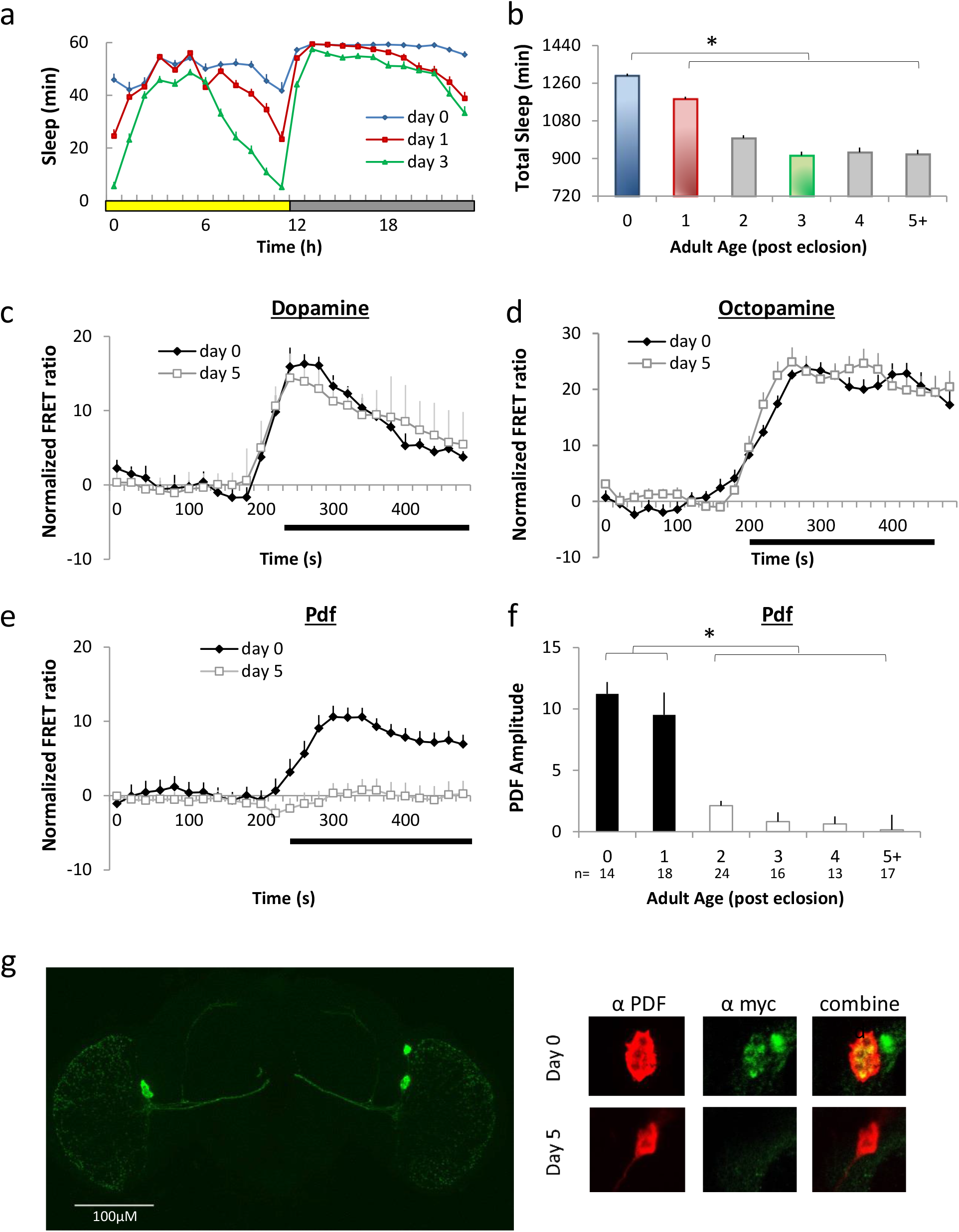
PDFR is expressed in lLNvs of young flies. **(a-b)** Sleep is elevated in young male flies following eclosion and reaches stable adult values in 3-d old flies (n=92-93 flies/age; One-way ANOVA for age, p=3.7^E-63^). **(c-e)** FRET ratio measurements in *Pdf-GAL4>UAS-Epac1* flies in response to Dopamine (3e-3M), Octopamine (3e-3M) and PDF (1e-6M) (n=5-15neurons/age). **(f)** The amplitude of lLNv responses to PDF decreases with age (PDF amplitude); (n=13-24neurons/age One-way ANOVA for age p=3.3^E-13^). **(g)** left: GFP expression in LNv neurons *(Pdf-GAL4;* UAS-*gfp*). right: Immunohistochemistry reveals co-expression of PDF (red) and myc (green) in 0 day old *P[acman] Pdfr-myc70* flies which is not observed on day 5. *p<0.05, modified Bonferroni test

### Expression of PDFR in lLNv neurons alters behavior in young flies

During the day, sleep is highest during the mid-day siesta and is reduced in the hours preceding lights out (Fig. 1a)^11,22,23^. We have operationally defined the 2 h period before lights out as the wake-maintenance zone based upon the observation that sleep rebound is absent or dramatically reduced when flies are released into recovery during this time window^22,27^. The ability to maintain waking in the face of high sleep-drive suggests that this window of time is protected for important waking behaviors^28^. With that in mind, we hypothesized that flies lacking the PDFR would sleep more than genetic controls during the wake-maintenance zone. *Pdfr*-null mutant *(Pdfr^5304^*) flies were outcrossed to *Cs* flies for 5 generations. To avoid handling of flies on the day they eclosed, *Pdfr^5304^* and *Cs* flies were plated on juice plates for 4 hours to lay eggs, and then L1 larvae were put into individual glass tubes and monitored. Sleep was assessed in male flies that eclosed between ZT1-ZT4. As seen in Fig. 2a,c on the day of eclosion *Pdfr^5304^* null-mutants sleep significantly more than their genetic controls during the wake-maintenance zone. To determine whether the change in sleep was due to expression of the PDFR in the lLNvs, we expressed wild-type *Pdfr* (*UAS-Pdfr^wt^*) using the *c929-GAL4* driver in a *Pdfr^5304^* mutant background. Since *c929-GAL4* is expressed in other peptidergic neurons^29^ we combined *c929-GAL4* with *cry-Gal80*, which targets the *GAL4* inhibitor *GAL80* to all clock neurons^30^. As seen in Fig. 2b,d sleep remained elevated during the wake-maintenance zone in *Pdfr^5304^;c929/+;cryGAL80/+* (green) and *Pdfr^5304^;UAS-Pdfr^wt^/+* (purple) parental controls as expected. In contrast, waking was rescued during the wake-maintenance zone in *Pdfr^5304^;c929/UAS-Pdfr^wt^* flies (red) and this increase in waking was prevented when the expression of *UAS-Pdfr^wt^* was blocked in clock cells *(Pdfr^5304^;c929/ UAS-Pdfr^wt^;cryGAL80/+*, blue). We verified the effectiveness of *cry-GAL80* using a *UAS-GFP* reporter (Supplementary Fig. 2a). To further exclude the possibility that expression of *UAS-Pdfr^wt^* in other peptidergic neurons outside the lLNvs altered waking, we rescued the expression *UAS-Pdfr^wt^* in a *Pdfr^5304^* mutant background using *Pdf-GAL4* which targets only LNv neurons. As seen in Supplementary Fig. 2b, sleep was reduced in *Pdfr^5304^;Pdf-GAL4/UAS-Pdfr^wt^* compared to parental controls. Finally, we asked whether the inability of *Pdfr^5304^* mutants to stay awake during the wake maintenance zone was due to the absence of *Pdfr* in the *lLNvs.* As seen in Fig. 2e, *Dcr2;929-GAL4/UAS-Pdfr^RNAi^* flies slept significantly longer during the wake-maintenance zone than either *Dcr2;c929-GAL4/+* or *UAS-Pdfr^RNAi^/+* parental controls. Together, these data indicate that PDFR in the lLNv promotes waking in young flies when sleep-drive is high.

**Fig 2.**
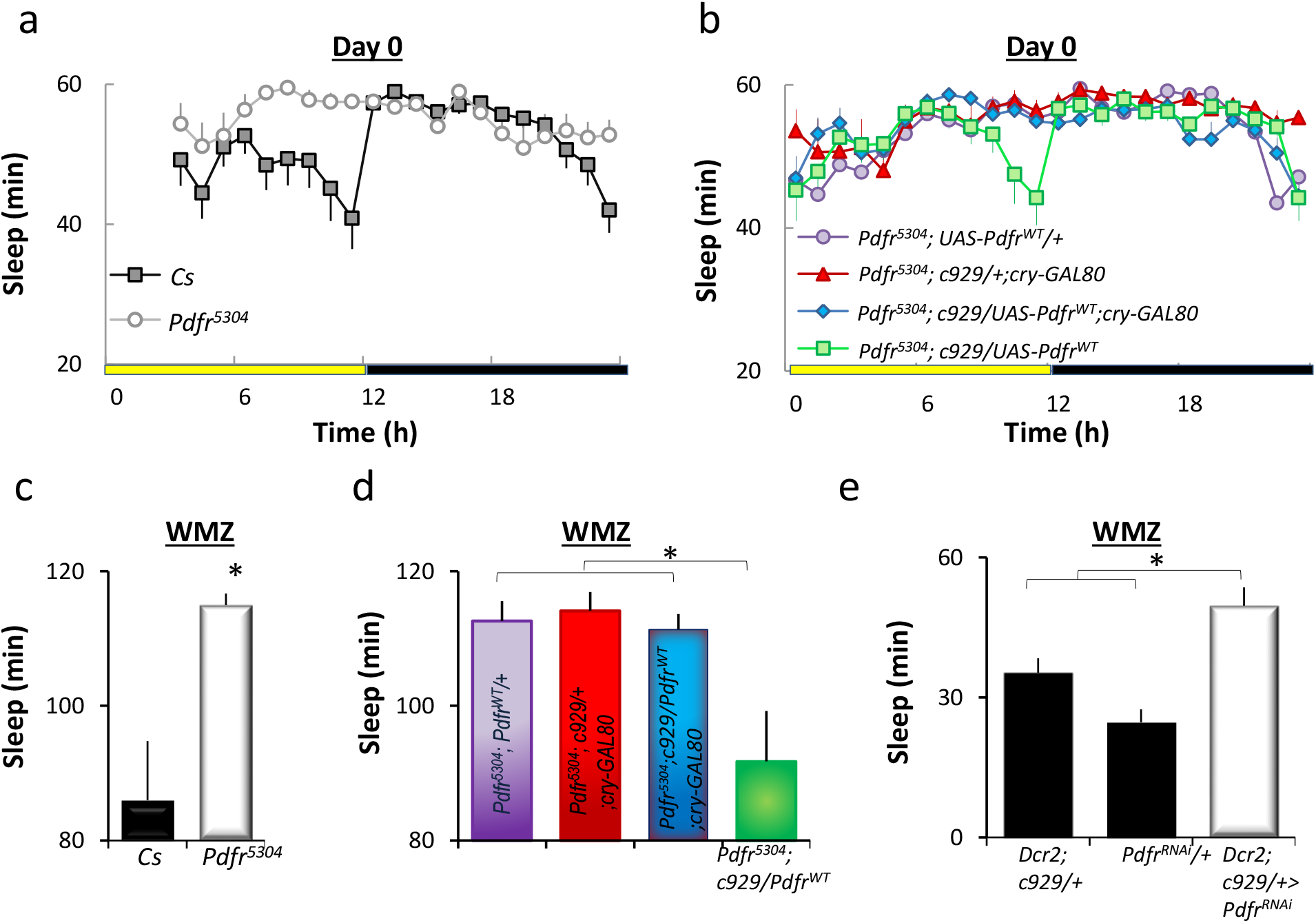
Expression of PDFR in lLNv neurons regulates sleep in young flies. **(a)** Sleep traces of *Pdfr^5504^* mutants and *Cs* controls on Day 0. **(b)** Sleep traces for *Pdfr^5504^; c929-GAL4; UAS-Pdfr* (rescue, green), *Pdfr^5504^; UAS-Pdfr/+, Pdfr^5504^; c929-GAL4/+; Cry-Gal80/+*, and *Pdfr^5504^;* c929-GAL4/*UAS-Pdfr*; *Cry-*Gal80 (n=26-31/genotype). **(c)** Quantification of sleep during the WMZ of flies shown in a. *Cs* flies sleep less during the WMZ than *Pdfr^5504^* mutants (n=26/genotype; t-test, p<0.05); **(d)** Quantification of sleep during the WMZ of flies shown in b. *Pdfr^5504^; c929-GAL4; UAS-Pdfr* sleep less than parental controls; ANOVA for genotype p=5.4^E-4^; n=22-31. **(e)** Sleep is increased in *Dcr2; c929-GAL4/UAS-Pdfr^RNAi^* flies on day 0 compared to *Dcr2; c929-GAL4/+* and *UAS-Pdfr/+* parental controls (ANOVA; p=1.04^E-5^; n=26-28).

Although the respecification of the PDFR in lLNvs supports waking in young flies, it is unclear whether the observed changes impact ecologically relevant behaviors. Inspired by the observation that the male pectoral sandpipers that sleep the least during breeding season sire more offspring^5^ we assayed mating success in flies with and without PDFR. As above, we began by evaluating *Pdfr^5304^* mutants and their genetic controls (*Cs*). Following eclosion, male flies were individually paired with a wildtype virgin female fly at ZT4 for 20 h and the pairings that produces offspring was tabulated. As seen in Fig. 3a, ~80% of pairings w/ Cs males resulted in offspring while only 25% of pairings with *Pdfr^5304^* mutants were successful on day 0. Moreover, mating success was also reduced when Pdfr was knocked down in c929-GAL4 expressing cells (Fig 3b). Importantly, the deficit in mating success observed in *Pdfr^5304^* mutants on day 0 was rescued by expressing wild-type PDF using c929-GAL4 (Fig. 3c). Previous studies have shown that the expression of PDFR in the dorsal lateral (LNd) neurons, a different set of clock neurons, promotes mating behavior in mature males^31^. However, no changes in mating success were observed in 2-d old *Pdfr^5304^* mutants or in *Pdfr^5304^;c929/UAS-Pdfr^wt^* rescue flies compared to genetic controls (Fig. 3a-c).

**Fig. 3.**
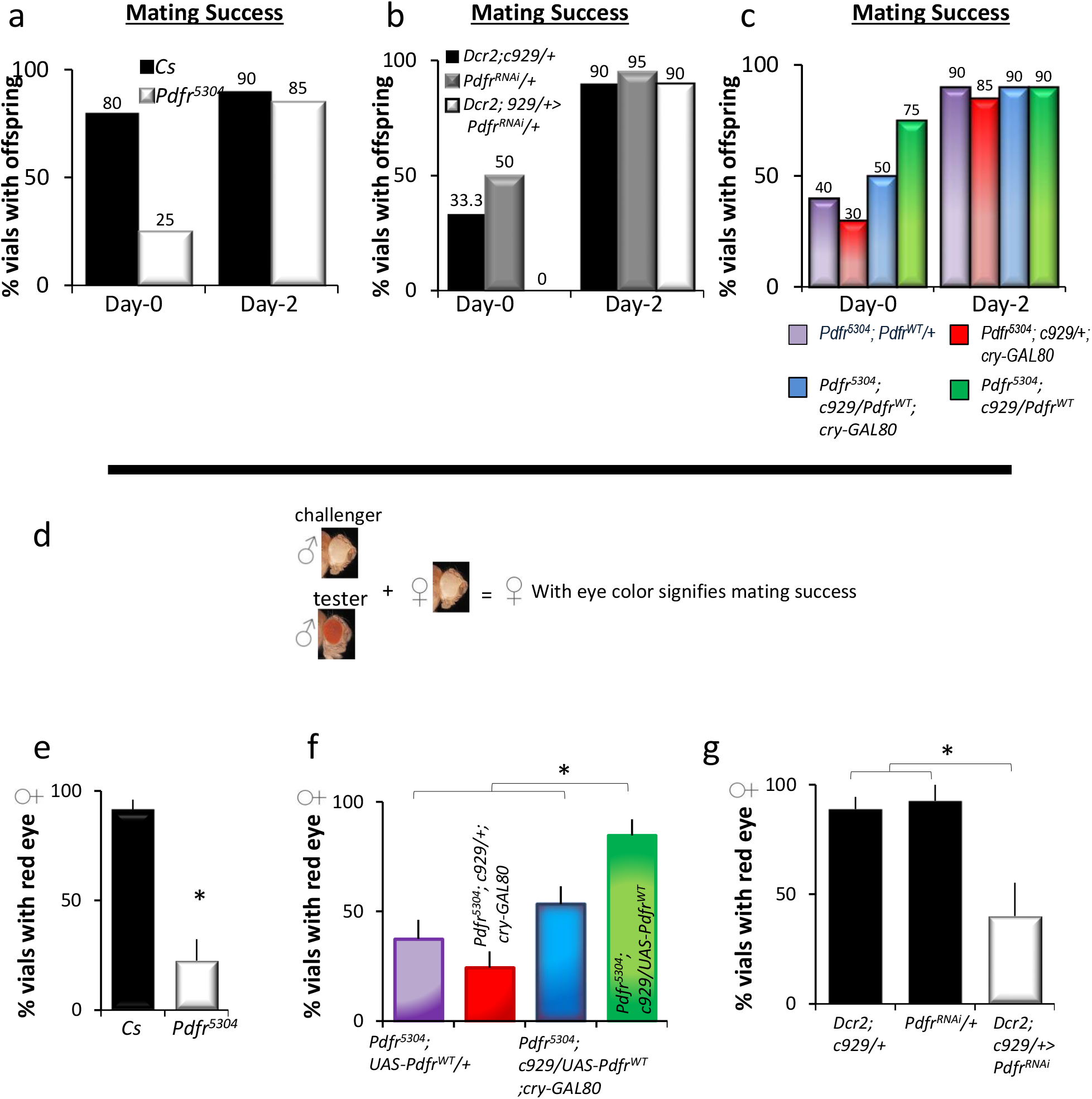
Role for PDFR in lLNv neurons in Mating Success. **(a)** *Cs* flies produce more offspring than *Pdfr^5504^* mutants (n=30/genotype). **(b)** *Dcr2; c929-*GAL4/UAS-*Pdfr^RNAi^* flies produce fewer vials with offspring compared to *Dcr2; c929-GAL4/+* and +/UAS-*Pdfr*^RNAi^ parental controls. **(c)** *Pdfr^5504^; c92-GAL4; UAS-Pdfr^WT^* (rescue, green) flies produce more offspring than *Pdfr^5504^; UAS-Pdfr^WT^/+* (purple), *Pdfr^5504^; c929-GAL4/+; CryGal80/+* (red), and *Pdfr^5504^; c929-*GAL4/UAS-*Pdfr*; *Cry-*Gal80 (blue) parental controls (n=30/genotype).*ANOVA, P<0.001. **(d)** Mating competition assay scheme on Day 1. **(e)** Cs males outcompeted white eye challenger flies compared to *Pdfr^5504^* mutants (t-test,P<0.001, n=60/genotype). **(f)** *Pdfr^5504^; c929-GAL4/+; UAS-Pdfr/+* males outcompeted white eyed challengers compared to *Pdfr^5504^; UAS-Pdfr, Pdfr^5504^/+; c929-GAL4/+; Cry-Gal80/+*, or *Pdfr^5504^; c929-GAL4/UAS-Pdfr; Cry-Gal80/+* controls (ANOVA p=4.9^−4^; n=3 sets of 20 flies/ line). **(g)** *Dcr2; c929-GAL4/UAS-Pdfr* RNAi flies displayed reduced mating success compared to *Dcr2; c929-GAL4/+* and *+/UAS-Pdfr* RNAi control flies (ANOVA, p=0.019, n=3 sets of 20 each line). *p<0.05, modified Bonferroni test.

To further determine whether expression of PDFR in the lLNvs was important for mating success we utilized a competition assay in which we rescued PDFR in a *Pdfr^5304^* mutant background. In this assay, one red-eye male and one white eyed male were combined with a white eye female for 2 h at the beginning of the wake-maintenance zone at zeitgeber time (ZT10) on Day 1. Successful mating of the red eye male was determined by female progeny with eye color (Fig. 3d). Consistent with the data presented above, *Pdfr^5304^* males sired fewer offspring than the Cs controls (Fig. 3e). Despite the fact that *white^−^* flies show impaired courtship^32,33^, white eyed males sired more offspring than red-eyed *Pdfr^5304^;c929/+, Pdfr^5304^; UAS-Pdfr^wt^*, and *Pdfr^5304^;c929/UAS-Pdfr^wt^;cryGAL80/+* controls (Fig. 3f). In contrast, male flies expressing the Pdfr in lLNvs *(Pdfr^5304^;c929/UAS-Pdfr^wt^)* sired more red eyed progeny on Day 1 (Fig. 3f). To determine whether the deficit in mating success in *Pdfr^5304^* mutants was due to loss of PDFR in the lLNvs, we evaluated *Dcr2;929-GAL4/UAS-Pdfr^RNAi^* flies. As seen in Fig. 3g, *Dcr2; c929>; UAS-Pdfr^RNAi^* lines reduced mating success compared to *Dcr2; c929/+* and UAS-*Pdfr*^RNAi^/+ parental controls. Therefore, the expression of the PDF receptor in the lLNvs is associated with successful mating in early adulthood when sleep pressure is high.

### Respecification of PDFR in ILNVs modulates adult behavior

Given that the expression of the PDFR in the lLNvs confers advantages to the young fly, we wondered why its expression would then be turned off on day 2-3 of adult life. To gain further insight into this question, we evaluated behavior in 5-d old flies ectopically expressing the PDFR in the lLNv using a specific split-GAL4 driver *(GRSS000645, lLNv-GAL4).* Daytime sleep was modestly reduced in *lLNv-GAL4>UAS-Pdfr^wt^* flies compared to *lLNv-GAL4/+* and *UAS-Pdfr^wt^/+* parental controls (Fig. 4a). As a negative control, we evaluated sleep in adult flies while expressing *UAS-Pdfr^RNAi^* in the lLNvs. Not surprisingly, expressing *UAS-Pdfr^RNAi^* in the lLNvs did not alter sleep in adult flies (Fig. 4b). Previous studies have shown that mutations that confer resistance in one environmental setting may increase the vulnerability of individuals in alternate settings^34^. Thus, we hypothesized that increased waking could sufficiently alter energy demands to make adult flies expressing PDFR in the lLNvs vulnerable to starvation. To test this hypothesis we starved flies and examined survival. As above, we examine the impact of starvation when the PDFR was overexpressed or knocked down in the lLNvs. As seen in Fig. 4c, survival was shorter in *lLNv>UAS-Pdfr^wt^* compared to *lLNv/ +* and UAS-*Pdfr*^wt^/+ parental controls. Astonishingly, *lLNv-GAL4>UAS-Pdfr^RNAi^* flies showed improved survival compared to both parental controls flies (Fig. 4d).

**Fig. 4.**
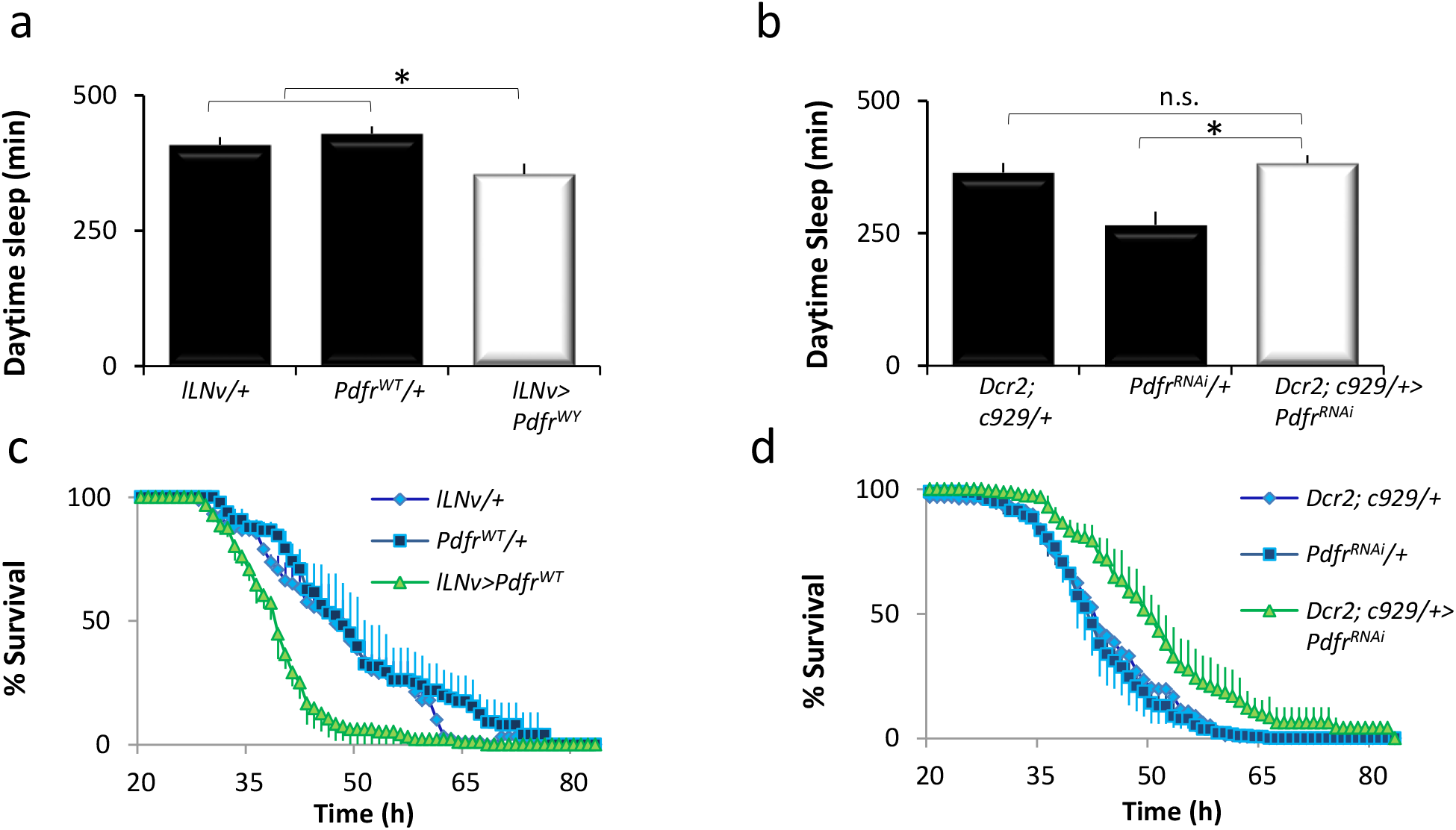
Behavioral Consequences of PDFR expression in lLNv neurons. **(a)** Daytime sleep in 5-day old *lLNv-GAL4>; UAS-Pdfr^WT^/+* flies and parental controls (ANOVA, p=0.004; n=30-32/genotype). **(b)** Sleep in *Dcr2; c929-GAL4/UAS-Pdfr* flies and parental controls (ANOVA p=1.86^E-07^; n=40-60/genotype; *p<0.05, modified Bonferroni test). **(c)** % survival during starvation in *lLNv-GAL4>; UAS-Pdfr^WT^* flies and parental controls (n=3 replicates of 10-16/genotype). **(d)** % survival during starvation in *Dcr2; c929 GAL4/UAS-Pdfr^RNAi^* flies and parental controls (n=3 replicates of 10-16/genotype).

The increased survival seen in starved *lLNv-GAL4>UAS-Pdfr^RNAi^* flies suggested that the genetic program that activates the PDFR in the lLNvs may be reactivated in mature adults during conditions of high sleep-drive. Short-periods of starvation (~12 h) increase waking without activating sleep-drive presumably to maintain cognition during foraging^14,34^. However, longer periods of starvation (~20 h) are able to activate homeostatic mechanisms^7^. Thus, we hypothesized that starvation would lead to the respecification of the PDFR in the lLNvs. As seen in Fig. 5a, PDF responses in the lLNvs of mature adult flies are restored following starvation when compared to age-matched, non-starved siblings. To determine how much time was required for starvation to respecify the PDFR in the lLNvs, we evaluated the time course of PDFR respecification in the lLNvs. Interestingly, starvation-induced restoration of PDF sensitivity in lLNvs requires a similar duration as reported for the activation of homeostatic drive (Supplementary Fig. 3a.). These data suggest that the respecification of the PDFR in the lLNvs is to help flies maintain wakefulness during starvation. With that in mind, we hypothesized that blocking the expression of the PDFR in the lLNvs would result in more sleep during starvation. Indeed, *Dcr2; c929>; UAS-Pdfr^RNAi^* flies slept more than parental controls between h 21 of starvation, when homeostatic drive begins, and 32 h of starvation prior to the point when flies begin dying (Supplementary Fig. 3b). In summary, starvation reduces sleep resulting in a build-up of sleep pressure which may mimic the conditions present in early adulthood that lead to PDFR respecification in lLNvs.

**Fig. 5.**
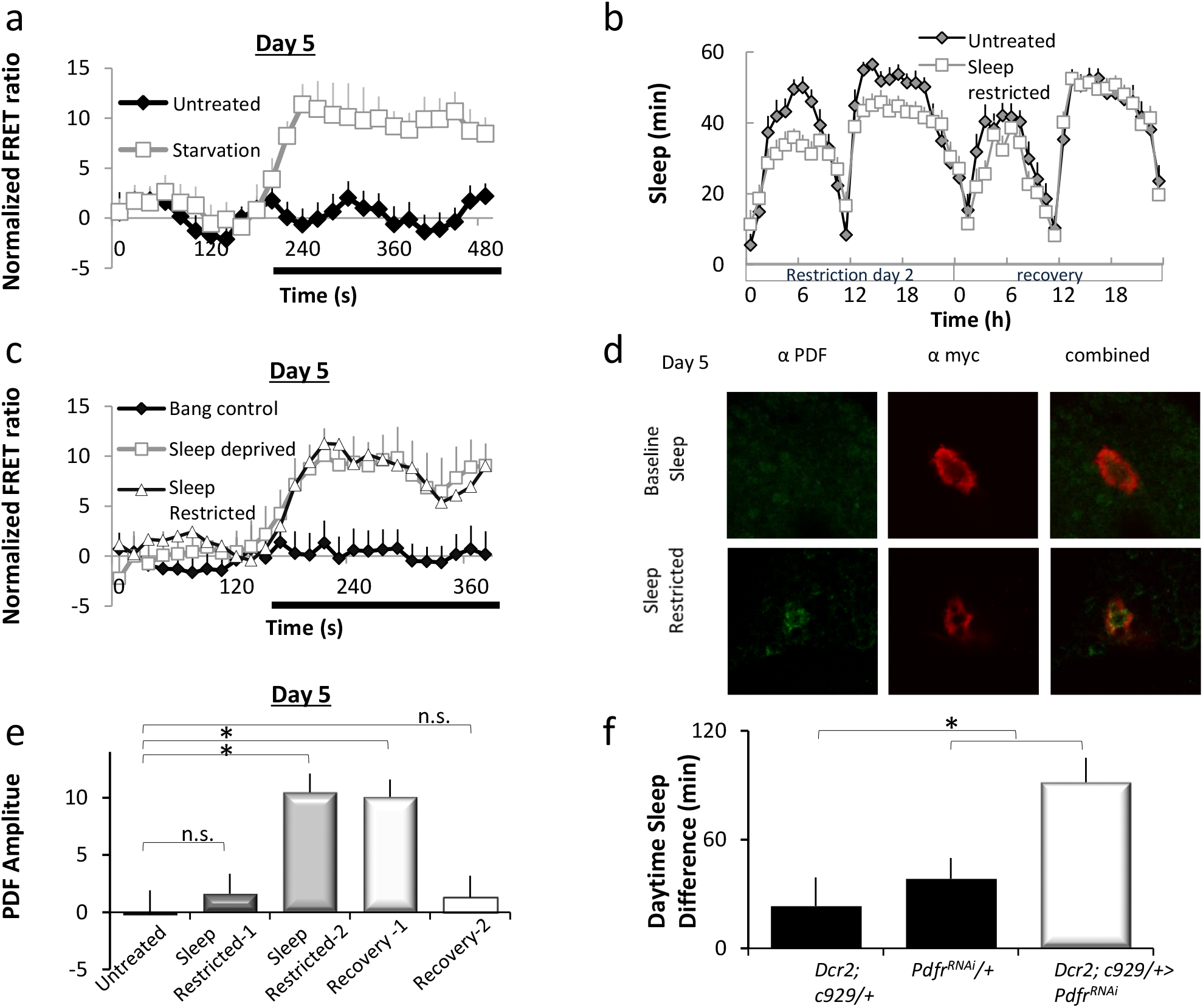
Sleep pressure-induces PDFR expression in mature lLNv neurons. **(a)** Normalized FRET ratio during PDF application in lLNV neurons from starved (n=10) and fed (n=16) *Pdf-GAL4>UAS-Epac1* flies. **(b)** Sleep in *Cs* flies on sleep-restriction day 2 and recovery. **(c)** Normalized FRET ratio during PDF application in lLNV neurons recorded from *Pdf-GAL4>UAS-Epac1* flies during sleep-restriction sleep-deprivation and bang controls (n=12-25neurons/genotype). **(d)** Immunohistochemistry of PDF (red) and myc (green) in 5-day old sleep restricted *P[acman] Pdfr-myc70* flies. **(e)** The amplitude of lLNv responses to PDF in lLNv neurons during baseline, sleep restriction and recovery (ANOVA; p=1.94^E-4^, n=9-24 neurons/condition). **(f)** Sleep rebound in *Dcr2; c929 GAL4/UAS-Pdfr^RNAi^* flies and parental controls (ANOVA p=2.22^E-3^; n=43-55). * p<0.05, modified Bonferroni test.

Starvation is an indirect method to increase sleep pressure. With that in mind, we asked whether sleep deprivation would also result in the respecification of the PDFR in the lLNvs in mature, adult flies. As seen in Fig. 5c, lLNvs respond physiologically to PDF following sleep deprivation in 5-d old flies. Although total sleep deprivation is the most common method for increasing sleep-drive in the laboratory, it seems unlikely that circumstances in the natural environment would keep an animal awake continuously for 12 h or more. In contrast, sleep consolidation is more easily disrupted and, perhaps, more likely to be impacted by a variety of environmental conditions^6,8^ Thus, we hypothesized that interrupting sleep consolidation would be sufficient to respecify the PDFR in the lLNvs. A variety of manipulations that increase sleep-drive (e.g. memory consolidation, activating the dorsal Fan Shaped body, etc.), increase average daytime sleep bout duration to >22 min/bout^35,36^. Thus we disrupted sleep consolidation by presenting a mechanical stimulus to the flies for 1 minute every 15 minutes for 48 h. As seen in Fig. 5b, this protocol modestly disrupted sleep and did not result in a compensatory sleep rebound. To determine if the lack of a sleep rebound was due to the respecification of PDFR, we examined the lLNvs physiologically and histologically. As seen in Fig. 5c, PDF responses in the lLNvs of mature adult flies are restored following 48 h of sleep restriction. To determine if the mechanical stimulus alone would respecify the PDFR in the lLNvs, siblings were exposed the same amount of stimulation (~190 minutes) as sleep restricted siblings but during the biological day when sleep debt does not accrue^11,37^. As expected, mechanical stimulation in the absence of sleep restriction did not respecify the PDFR in the lLNvs (Fig. 5c). To confirm the physiological data, PDFR was examined directly using *Pdfr-myc*^26^. MYC antibody staining in the lLNvs is clearly visible in mature adult flies following sleep restriction but is not observed in non-disturbed age-matched controls (Fig. 5d). Next we asked how much sleep restriction was required for the respecification of the PDFR. As seen in Fig. 5e, PDF sensitivity becomes apparent after 24-48 hrs of sleep restriction, is sustained during the first day of recovery, and then dissipates. Finally, we asked whether knocking down the PDFR in the lLNvs would modulate sleep homeostasis following sleep disruption. As seen in Fig. 5f, *Dcr2;c929/+>UAS-Pdfr^RNAi^* flies slept significantly more following sleep restriction than *Dcr2;c929/+ and UAS-Pdfr^RNAi^/+* parental controls. These data indicate that PDFR can be respecified to mitigate against the effects of sleep pressure in the context of sleep disruption.

### nejire modulates PDFR in both young and mature lLNv neurons

The PDFR is transiently expressed in the lLNvs of young flies and can be respecified again in mature adults in response to certain environmental perturbations. Thus, we asked whether these seemingly different conditions invoke the same mechanisms to activate the expression of PDFR in the lLNvs. To begin, we conducted an RNAi screen of transcription factors that are known to be expressed in LNvs^38^. We crossed UAS-RNAi lines with *pdf-GAL4;UAS-Epac* and monitored PDF sensitivity in both lLNvs and sLNvs in young flies on day 0. As mentioned above, sLNvs display persistent expression of the PDFR in both young and mature-adults. Thus, we hypothesized that by monitoring both cell types, we could distinguish between regulatory elements specific to the transient pathway in lLNvs. We also examined dopamine responses to discriminate between transcription factors specifically involved in the PDF pathway and those common to other GPCR signaling pathways. As seen in Fig. 6a, knocking down *Drosophila CREB-binding-protein (nejire)* or *Suppressor of Under-Replication (SuUR)* ablated PDF sensitivity in lLNvs on Day 0 while other transcription factor left the sensitivity of the lLNvs to PDFR largely intact. The amplitude of DA responses was not altered by *nejire, SuUR*, or any other RNAi lines revealing the roles of *nejire* and *SuUR* are specific to the PDF pathway in this context (Supplementary Fig. 4a). PDF sensitivity in the sLNvs was not ablated by RNAi knockdown of *nejire* (Supplementary Fig. 4b). Interestingly, *nejire* also plays a role in the respecification of the PDFR in the lLNvs in mature-adults following sleep restriction (Fig. 6b). As in young flies, the panel of RNAi lines did not alter DA responses in the lLNvs (Supplementary Fig. 4c). To further evaluate the role of *nejire* in the respecification of the PDFR in mature adults, we expressed wild-type *nejire (UAS-nejire^WT^)* or *UAS-nejire^RNAi^* using *Pdf-GA4; UAS-Epac.* We hypothesized that the overexpression of *nejire* would restore PDFR sensitivity to the lLNvs in well-rested mature-adults and that knocking down *nejire* would block the respecification of the PDFR in the lLNvs during sleep restriction. Indeed, the sensitivity of the lLNvs to PDF was restored in well-rested mature-adults by overexpressing *UAS-nejire^WT^*. Conversely, the respecification of the PDFR to the lLNvs during sleep restriction was blocked by *UAS-nejire^RNAi^* (Fig. 6d). Together these data reveal that conditional PDFR expression in lLNvs shares common mechanisms in both young flies and mature-adults.

**Fig. 6.**
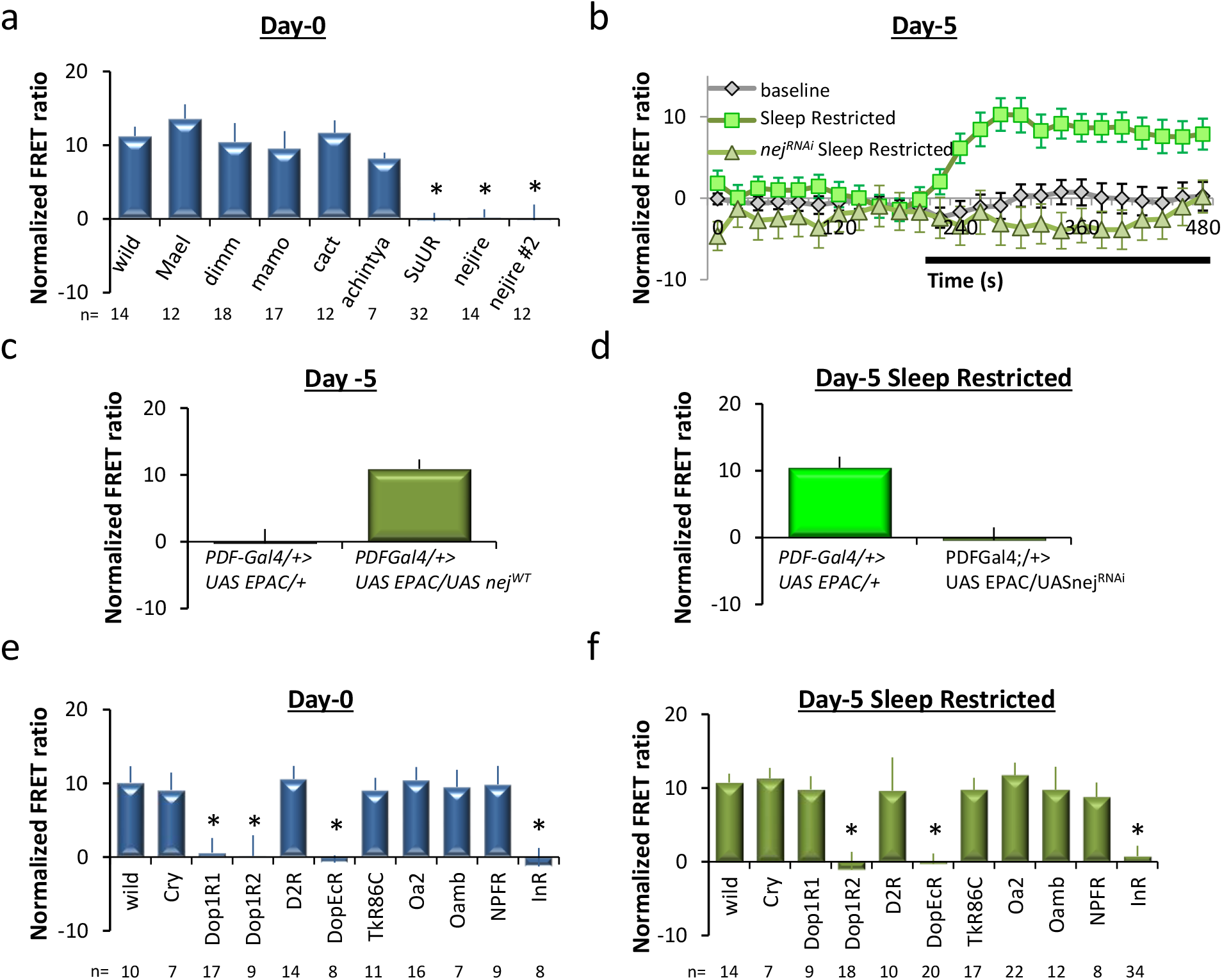
*nejire* regulates PDFR respecification in lLNvs of both young and mature flies. **(a)** The amplitude of lLNv responses to PDF on Day 0 in *Pdf-GAL4>UAS-Epac1* flies crossed to *UAS-RNAi* lines of the depicted transcription factors (ANOVA; p=1.26^E-11^, * p<0.05, modified Bonferroni test, n are listed below the x-axis). **(b)** PDF amplitude in control and 5-d old sleep restricted *Pdf-GAL4>UAS-Epac1* flies compared to *Pdf-GAL4>UAS-Epac1* flies expressing *UAS-nej^RNAi^* (n=6-19 neurons/genotype). **(c)** In the absence of sleep loss, the lLNvs of *Pdf-GAL4>UAS-Epac1/UAS-nej^WT^* respond to PDF while age matched *Pdf-GAL4>UAS-Epac1* do not (t-test, P<0.001, n=9-26 neurons/genotype). **(d)** Data quantified from b. (t-test, p=0.001, n=16-26). **(e)** The amplitude of lLNv responses to PDF in lLNv neurons on Day 0 in *Pdf-GAL4>UAS-Epac1* flies crossed to *UAS-RNAi* lines of the depicted cell surface receptors (ANOVA p=2.8^−6^ p<0.05, modified Bonferroni test, n are listed below the x-axis). **(f)** The amplitude of lLNv responses to PDF in lLNv neurons on Day 5 in sleep restricted *Pdf-GAL4>UAS-Epac1* flies crossed to *UAS-RNAi* lines of the depicted cell surface receptors (ANOVA p=3.3^E-9^ p<0.05, modified Bonferroni test, n are listed below the x-axis).

Finally, we asked whether similar mechanisms were used by young and mature adults for the activation of *nejire.* To identify cell surface receptors that might interact with *nejire*, we once again consulted a database of genes known to be enriched in the LNvs^38^. We then conducted a targeted RNAi screen to evaluate PDF sensitivity in young flies and mature-adults following sleep restriction. Interestingly, both young flies and sleep-restricted adults, display reduced or absent responses to PDF when knocking down the *Dopamine/Ecdysteroid receptor (DopEcr), Dopamine 1-like receptor 2 (Dop1R2)*, and the *Insulin-like receptor (InR)* while other cell-surface receptors known to be expressed in the LNvs were without effect (Fig. 6e,f). Importantly, knocking down cell-surface receptors did not alter dopamine sensitivity in any fly tested (Ext Data fig. 4c, d). Together these data indicate that the respecification of the PDFR in the lLNvs involved the *DopEcR, Dop1R2*, and *InR* in both young flies and mature adults during sleep restriction.

## Discussion

A growing number of studies indicate that sleep regulatory mechanisms are plastic and can be harnessed to match an individual’s sleep need with environmental demands^6,14,39^ Although Hebbian and synaptic plasticity modulate circuit function in a variety of contexts, these forms of plasticity may not be well suited to sculpt the balance of sleep and wake-promoting circuits at specific times of day^40^. In contrast, receptor respecification is a form of plasticity that may allow an individual to engage in adaptive waking behaviors at optimal circadian times while still allowing the animals to obtain needed sleep at other times^3^. Indeed, our data indicate that the PDFR is transiently expressed in wake-promoting clock neurons during the first~48 h of adult life when sleep drive is high. The associated increase in waking is confined to a small portion of the circadian day and supports mating success and mating competition. In contrast, the response properties of the lLNvs to the global wake-promoting transmitters octopamine and dopamine remains unchanged^41^. Interestingly, when sleep is disrupted in 5-day old adults, the PDFR is once again expressed in the lLNvs. Thus, targeted receptor-respecification may be an effective strategy that can be used to support important, species-specific behaviors during conditions of high sleep drive without substantially disrupting the ability of the animal to obtain needed sleep.

Our data indicate that there is a strong relationship between sleep drive and the respecification of the PDFR in a subset of clock neurons. That is, while the lLNvs are unresponsive to PDF in mature adults^24^, the lLNvs display robust responses to PDF following sleep deprivation, sleep restriction and prolonged starvation. Importantly, no changes in the response properties of the lLNvs were observed when the animals were exposed to the mechanical stimulus in the absence of sleep restriction. Interestingly, the response properties of the lLNvs was not visible until the second day of sleep restriction indicating that low amounts of sleep drive are not sufficient to respecify the PDFR. Consistent with this hypothesis, short-durations of starvation induce episodes of waking that are not compensated by a sleep rebound^42^ and do not change the response properties of the lLNvs to PDF. In contrast, after ~20 of starvation, a time when flies begin to display a sleep rebound, the lLNvs begin to respond to PDF. These data suggest that the PDFR may be respecified in the lLNvs to, for example, help sleepy animals stay awake long enough to support a basal level of foraging. Indeed, knocking down the *Pdfr* in clock neurons results in a larger sleep rebound following sleep restriction. Increased sleep in many circumstances may be maladaptive since it would likely limit the opportunity to forage or mate^43^.Together these data support the hypothesis that the PDFR is expressed to assist waking behaviors during conditions of high sleep drive.

Given that high sleep drive can negatively impact male sexual behavior (9), it is curious that the PDFR is not typically expressed in the lLNvs of healthy adults. However, previous studies have shown that genes that confer resilience to specific environmental challenges can be deleterious in other circumstances (31). Indeed, the exogenous expression of PDFR in the lLNvs during adulthood reduced survival during prolonged starvation. These data suggest that the normal downregulation of PDFR expression in lLNv neurons of healthy adults may be advantageous in that it removes potentially excessive behavioral drives that could deplete valuable resources. Indeed, genetically preventing PDFR expression in lLNv neurons during starvation extended survival.

Although sleep drive does not change the response properties of the lLNvs to dopamine, our data suggest that changes in dopaminergic tone may play a role in the respecification of the PDFR in the lLNvs. Specifically, knocking down specific dopamine receptors in the lLNvs prevents the respecification of the PDFR in both young flies and sleep restricted 5-day old adults. Although the precise dopaminergic neurons have not yet been identified, the PPL2 dopaminergic neurons project to the ILNv’s to promote wakefulness^44^ and may play a role in the expression of the PDFR in lLNvs. In addition to dopamine receptors, our data identify a role of the transcription factor *nejire*, (cAMP-response element-binding protein), in promoting the expression of the PDFR during conditions of high sleep drive. Interestingly, *nejire* plays a role in circadian function where it has been suggested to allow cross-talk between circadian transcription and the transcriptional regulation of other important processes such as sleep, metabolism, and memory formation^45,46^.

Previous studies have shown that activity-dependent respecification of receptors in mammals can occur in adult neurons in response to >1 week of sustained increases in sensory activity^2,3^ The most common forms of respecification alter the polarity of the synapse to alter the function of the circuit^3^. Our data suggest an additional type of respecification in which an input pathway into a circuit can be turned on and off, without changing the sign of the synapse (excitatory/inhibitory). Presumably, turning on an input pathway may be a rapid first step to balance the impact of sustained activity in opposing circuits (e.g. sleep vs. wake). However, enhancing the activity of a circuit may create a positive-feedback loop which can destabilize the system and lead to adverse consequences. Indeed, while the respecification of the PDFR in the lLNvs can improve mating success during high sleep-drive, it also results in early lethality during starvation. Understanding how sleep-drive modulates respecification-plasticity in other sleep regulatory circuits may provide critical insight into the role that sleep plays in maintaining adaptive behavior in an ever changing environment.

## Method

### Flies

Flies were cultured at 25°C with 50-60% relative humidity and kept on a diet of yeast, dark corn syrup and agar. Newly-eclosed males were collected and entrained 4-7 days in a 12h:12h Light:Dark (LD) cycle, unless otherwise specified. RNAi stocks were obtained from VDRC and TRiP stock centers. *DopEcR^RNAiJF03415^, Dop1R1^RNAiHM04077^, Dop1R2^RNAiHMC06293^, D2R^RNAiHMC02988^, InR^RNAiHMS03166^, NPFR^RNAiJF01959^, Oamb^RNAiJF01673^, TkR89C^RNAiJF02160^, Oa2^RNAiHMJ22156^, Cry^RNAiJF01880^, Mael^RNAiHMS00102^, dimm^RNAiHMS01742^, mamo^RNAiHMC03325^ cac^RNAiHM0402^, achintya^RNAiHMS01127^, SuUR^RNAiGL0108^, nejire^RNAihp12^, nejire^RNAihp123.3^*. Other stocks used were *c929(dimm)-GAL4; PDF-GAL4; lLNv^GRSS000645^-GAL4;, UAS-nejire^wt-V5^*. All other UAS lines and GAL4 lines have been described previously: *Pdfr*-null mutant (*Pdfr^5304^*); *UAS-Pdfr^wt^; w; UAS-Epac1camps50A (19), w, Pdf-GAL4(M)* and *UAS-Pdfr^RNAi^* (44). *c929-GAL4; cry-GAL80/UAS-GFP* flies and *P[acman] pdfr-myc70 flies* (22) were used for immunolabelling.

### Sleep

Sleep was measured as described previously^11^. In short, individual flies were placed into ~65 mm tubes which were then placed into Trikinetics Drosophila Activity Monitoring System (www.Trikinetics.com, Waltham, Ma). Locomotor activity was monitored using an infrared beam, and was assessed using 1-minute time bins. Sleep has been defined as periods of quiescence lasting 5 minutes or longer^11^.

### Mating Success

Mating success assay consisted of putting one virgin female and one male of varying genotype, either wildtype, null PDFR background, ectopically PDFR expressed in null background, or restored PDFR to lLNv neurons in null background. The pair of flies was put into a vial at ZT4 on day 0 of adulthood and then the male was removed at ZT24. Mating success was determined days later through visual inspection for offspring (pupae, larvae, etc.). Females from vials that produced no offspring were subsequently provided several males to test for her reproduction viability.

### Mating Competition

A mating competition assay was also carried out using two males who compete for one female. In each tube one white eye male, and one red eye male of varying PDFR levels competed to mate with a white eye female. The two competing males were added to a vial simultaneously with a mature virgin female, just prior to the wake maintenance zone (ZT10) and left in the vial until the end of WMZ (ZT12). Successful mating of the red eye male was determined by female progeny with eye color. Twenty competitions were set up for each genotype and repeated three times. Only competitions resulting in progeny were used for analysis.

### Sleep Restriction

Disruption of sleep was performed similarly as previously described^27,47^. Flies were placed into individual 65 mm tubes and a sleep-nullifying apparatus (SNAP) which mechanically disrupted sleep for one minute every fifteen minutes for 24-48 hours, which both reduced and fragmented sleep. Sleep homeostasis was calculated for each individual as a ratio of the minutes of sleep gained above baseline during the 48 h of recovery divided by the total min of sleep lost during 12 h of sleep deprivation.

### Physiology

Methods generally followed those of Klose et al., (2016). Flies were removed from DAM monitors and glass tubes were placed on ice for approximately 5 minutes. 3-4 flies were pinned onto a sylgaard dissection dish, and were dissected in cold calcium-free HL3 (Stewart et al., 1994). Dissected brains were transferred onto a poly-lysine treated dish (35 3 10 mm Falcon polystyrene) containing 3 ml of 1.5mM calcium HL3. Two to four brains were assayed concurrently, typically a mutant line and its genetic controls. Image capture and x,y,z stage movements were controlled using SLIDEBOOK 5.0 (Intelligent Imaging Innovations), which controlled a Prior H105Plan Power Stage through a Prior ProScanII. Multiple YFP/CFP ratio measurements were recorded in sequence from each hemi-segment of each brain in the dish. Following baseline measurements, 1 ml of saline containing various concentrations of either PDF, DA, or OA (Sigma-Aldrich) was added to the bath (dilution factor of 1/4). We tested normality in the data using the Shapiro-Wilk test in SigmaPlot (Systat Software), maximum amplitude values were used to perform ANOVA analyses followed by post hoc Tukey tests.

### Immunocytochemistry

Whole flies were fixed in 4% PFA for several hours (???), and brains were then dissected in ice cold PBS and incubated overnight with the following primary antibodies: mouse anti-PDF, (5F10, 1:10 dilution, Hybridoma Bank, University of Iowa???), chicken anti-myc (GFP-1020; 1:1000???), and anti-GFP. Secondary antibodies were Alexa 488 and 633 conjugated at a dilution 1:200. Brains were mounted on polylysine treated slides in Vectashield H-1000 mounting medium. Confocal stacks were acquired with a 0.5μm slice thickness using an Olympus FV1200 laser scanning confocal microscope and processed using ImageJ.

### Statistics

All comparisons were done using a Student’s T-test or, if appropriate, ANOVA and subsequent planned comparisons using modified Bonferroni test unless otherwise stated. All statistically different groups are defined as *P < 0.05.

### Author Contributions

M.K.K and P.J.S. designed the experiments and wrote the paper. M.K.K. and P.J.S. performed the experiments. M.K.K. and P.J.S. analyzed the data.

## Acknowledgments

We thank Gerald Rubin and Paul Taghert for sharing reagents and flies. We also thank Stefan Dissel for help throughout this project, Lijuan Cao for help with immunohistochemistry and Krishna Melnattur for comments. This study was funded by NIH Grants R01-NS051305-01A1, R01NS057105, to PJS and the NIH Neuroscience Blueprint Core Grant, #NS057105.

## Extended data Figure Legends

**Extended Data Figure 1.**
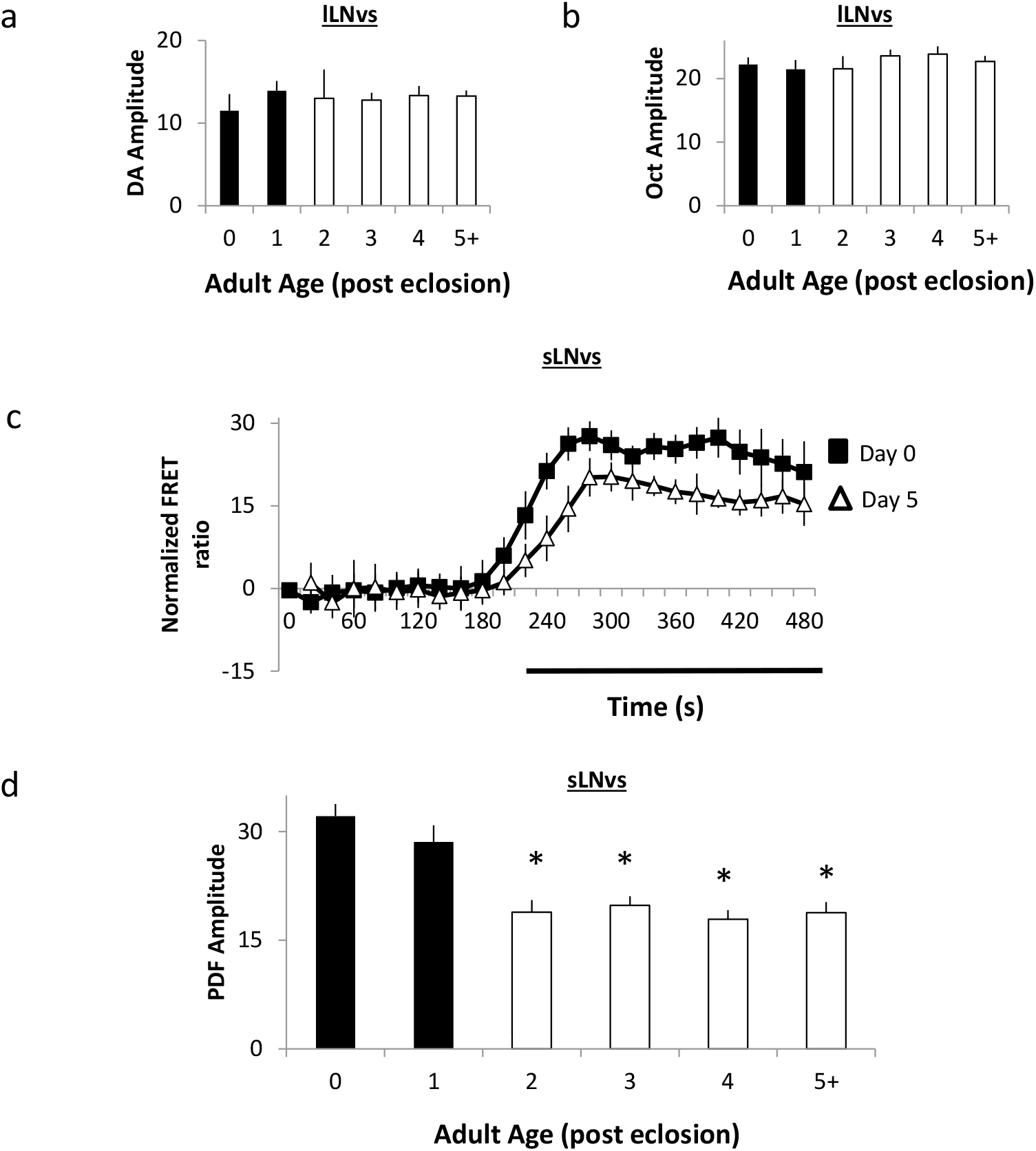
a-b. Response of lLNvs to Dopamine and Octopamine in *Pdf-GAL4>UAS-Epac1* flies from day 0 to day 5+ (n=4-15 neurons per age, ANOVA F_[5,36]_=6.08,p=0.96 and ANOVA F_[5,54]_=8.93,p=0.90 respectively). c. Normalized FRET ratio in small LNv neurons before and during PDF exposure on day 0 (n=6) and day 5 (n=7). d. PDF response amplitude in sLNv neurons on day 0 to day 5+ (ANOVA F_[5,76]_ = 13.31,p=3.75^E-9^ n=8-20 neurons per age). *p<0.05, modified Bonferroni test.

**Extended Data Figure 2.**
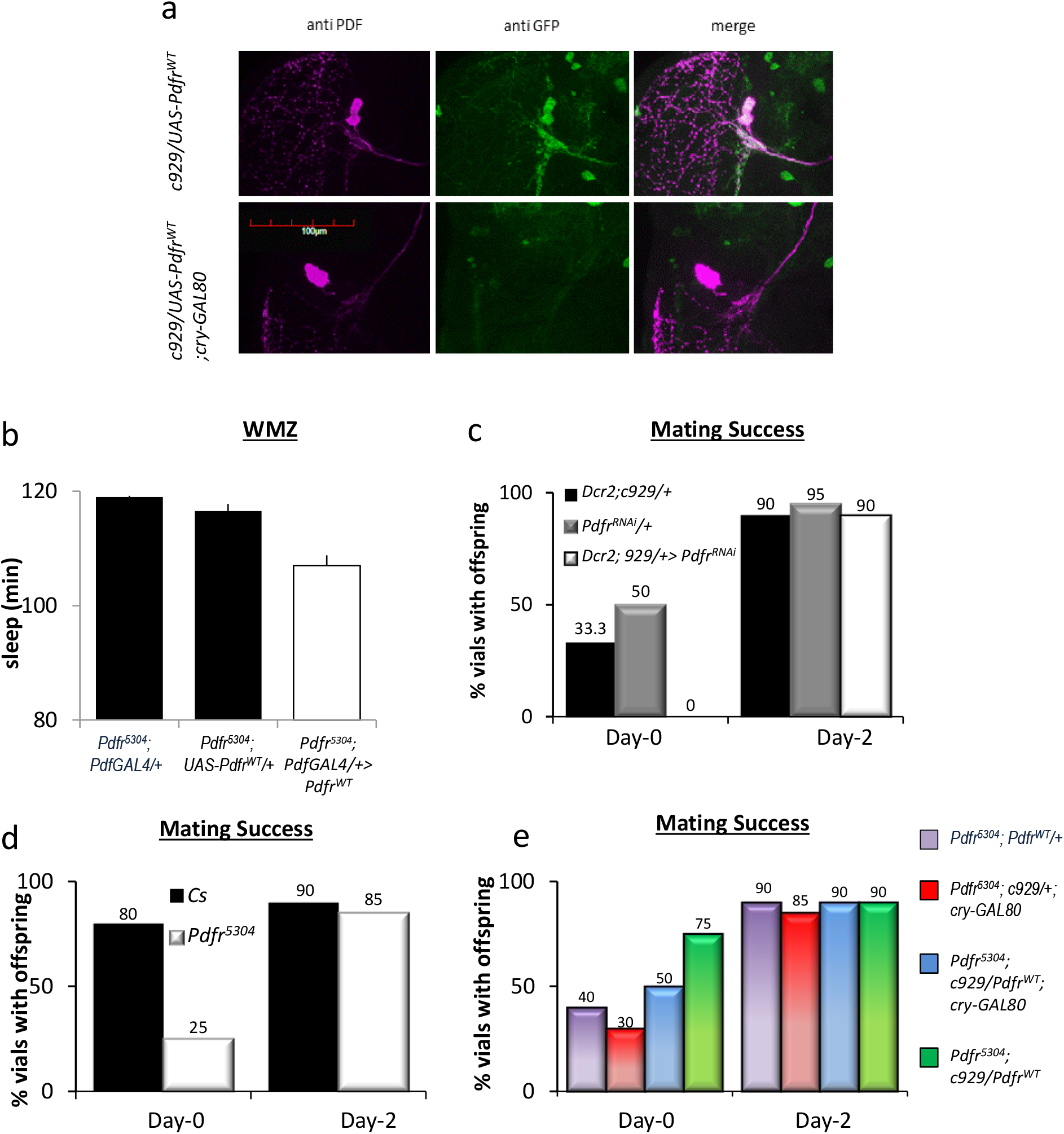
a. Immunohistochemistry for PDF and GFP reveals the expression of GFP in the lLNvs of *c929-GAL4/UAS-gfp* flies but not in the brains of *c929-GAL4/UAS-gfp; Cry-Gal80 flies.* b. *Pdfr^5504^; PDF>/UAS-Pdfr^WT^* flies exhibit more waking during the wake maintenance zone (WMZ) than *Pdfr^5304^; Pdf-GAL4/+* and *Pdfr^5304^;UAS-Pdfr^WT^/+* parental controls (ANOVA F_[2,91]_=4.63,p=0.01 n=41-64 flies/genotype) flies.

**Extended Data Figure 3.**
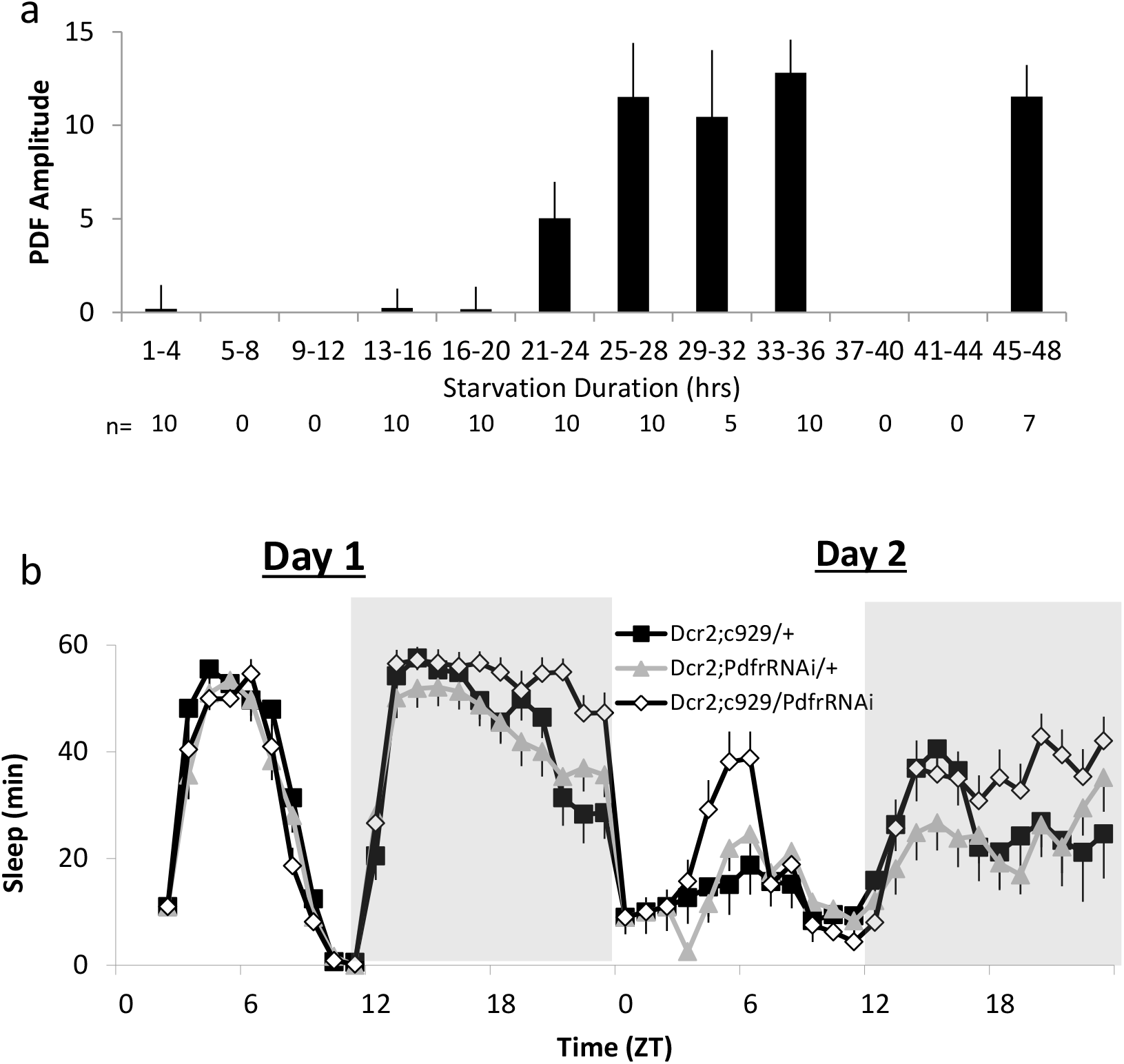
a. The amplitude of lLNv responses to PDF is observed in 5-day old *Pdf-GAL4>UAS-Epac1* flies following 21-24 hours of starvation. Data are shown for 4 h bins (ANOVA F_[7,71]_=9.08,p=9.8^E-8^; n is as indicated beneath each bin). ANOVA, P<0.001, see table. b. Sleep (minutes) during 48 hours of starvation in *Dcr2; c929 GAL4/UAS pdfr^RNAi^* flies (n=24), *Dcr2; c929-GAL4/+* (n=18) and +/UAS-*Pdfr^RN^A* (n=32) control flies.

**Extended Data Figure 4.**
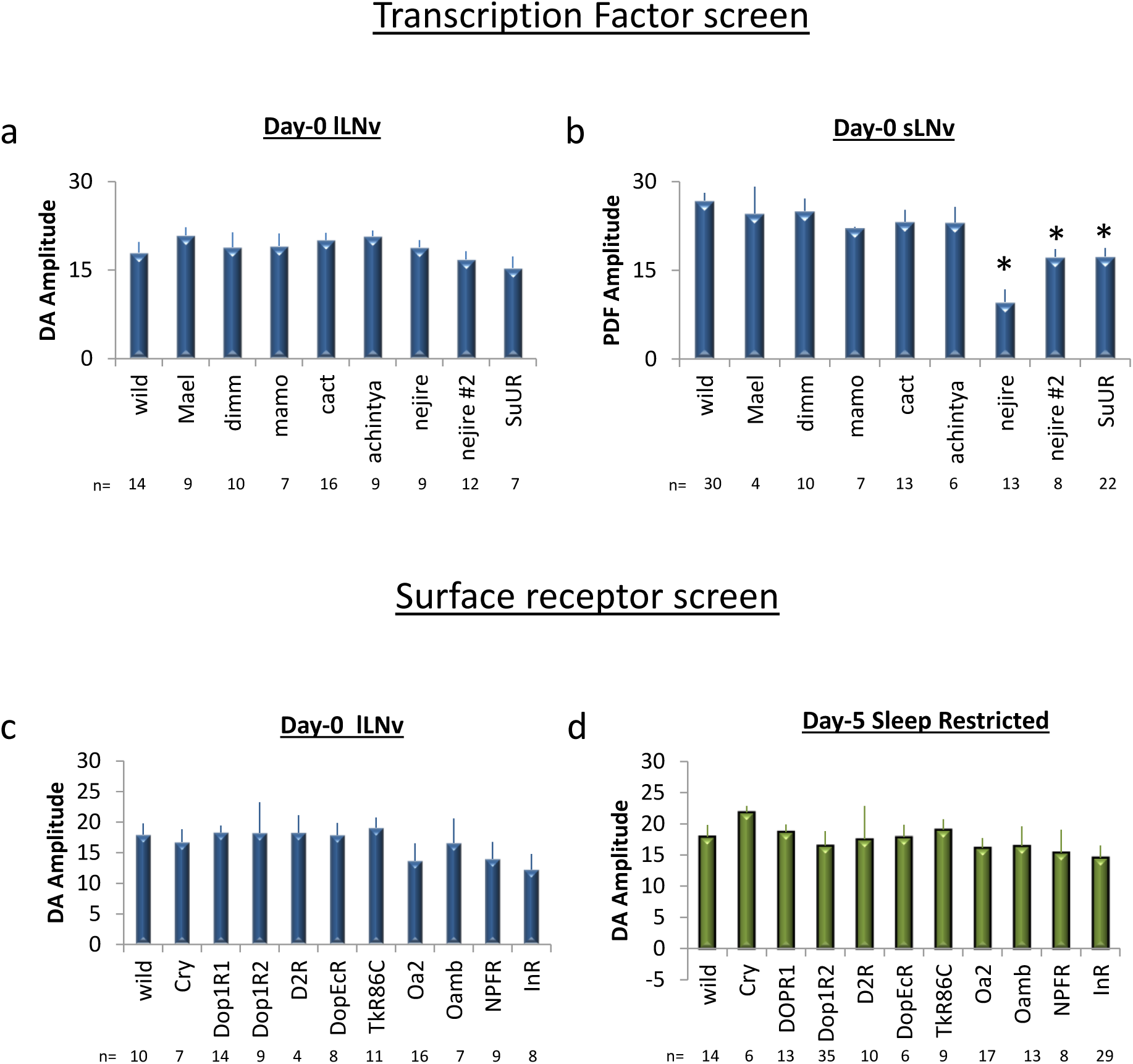
a. The amplitude of lLNv responses to Dopamine on day 0 in *Pdf-GAL4>UAS-Epac1* flies coexpressing RNAi lines for the depicted transcription factors (ANOVA F_[8,92]_=1.04,p=0.42; n is as indicated beneath each bin). b. The amplitude of sLNv neurons responses in *Pdf-GAL4>UAS-Epac1* flies co-expressing RNAi lines for the depicted transcription factors neurons on day 0(ANOVA F_[8,112]_=9.36,p=1.19^E-9^). c. The amplitude of sLNv neurons responses to Dopamine on day 0 in in *Pdf-GAL4>UAS-Epac1* flies co-expressing RNAi lines for the depicted cell surface receptors (ANOVA F_[10,108]_=0.79,p=0.63; n is as indicated beneath each bin). d. The amplitude of lLNv neurons responses to Dopamine following sleep restriction in 5-day old *Pdf-GAL4>UAS-Epac1* flies co-expressing RNAi lines for the depicted cell surface receptors (ANOVA F_[10,159]_=0.42,p=0.94 n is as indicated beneath each bin).

**Extended Data Table 1.**
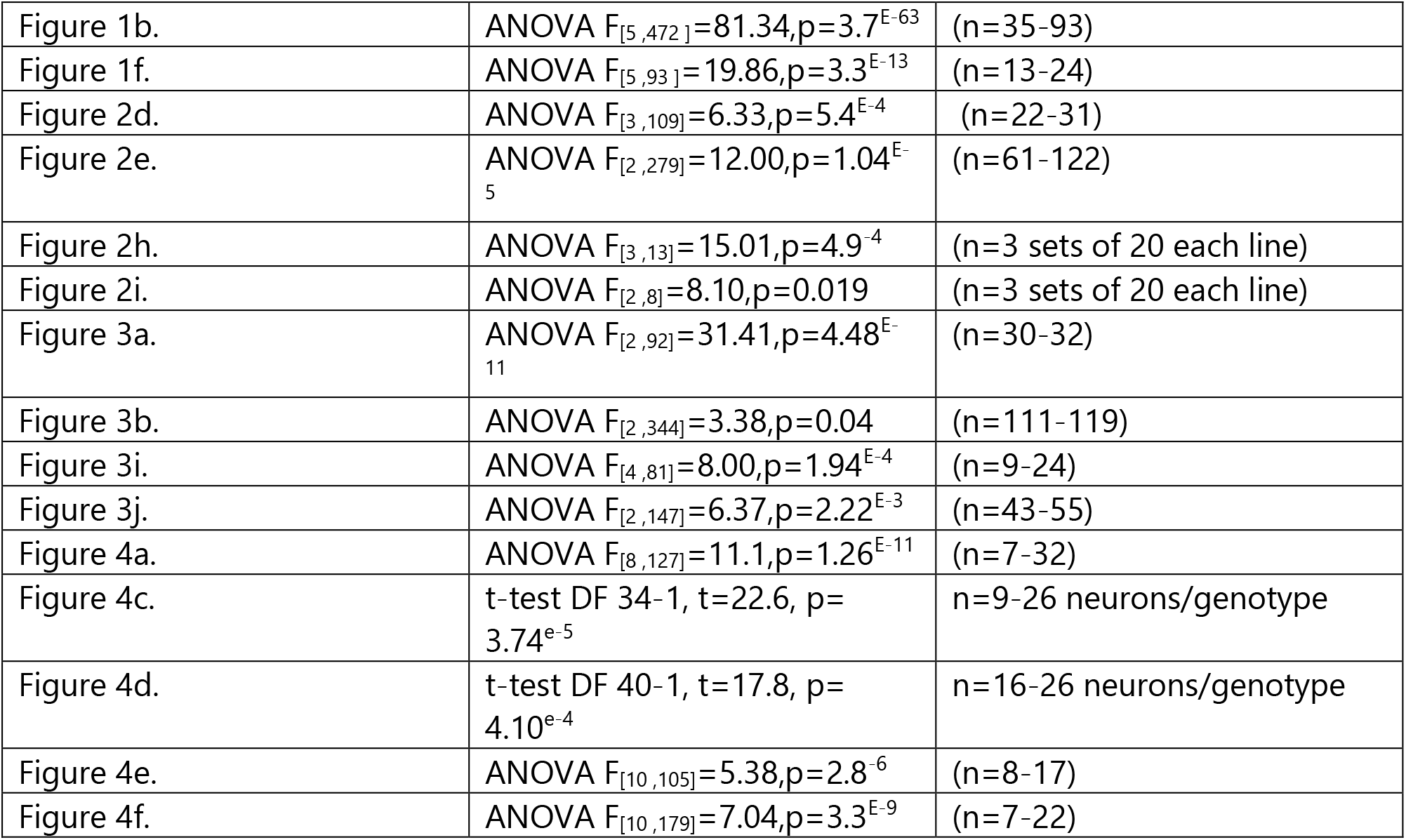
Statistics

